# Real-Time Neurotransmitter Sensing in Midbrain Organoids Using 3D Carbon Cup Electrodes

**DOI:** 10.64898/2026.10.12.758473

**Authors:** Samuel Tavares da Silva Maraschin, Surangrat Thongkorn, Swetha Vasudevan Kanakkottu, Babak Rezaei, Giorgio Scordo, Edoardo Sozzi, Malin Parmar, Arto Heiskanen, Stephan Sylvest Keller, Jenny Emnéus

**Affiliations:** DTU Bioengineering, Technical University of Denmark, 2800 Lyngby, Denmark; DTU Nanolab, Technical University of Denmark, 2800 Lyngby, Denmark; Developmental and Regenerative Neurobiology, Lund Stem Cell Center, Department of Experimental Medical Science, Faculty of Medicine, Lund University, Sweden; Center for Analysis and Synthesis, Department of Chemistry, Lund University, Sweden

## Abstract

Real-time, chemically selective detection of neurotransmitter release in brain organoids remains a challenge due to their complex three-dimensional architecture and limited optical accessibility. We introduce 3D-printed carbon cup electrodes (CCEs) -pyrolyzed, multi-level pillar structures - designed to confine a single organoid, increase electroactive surface area, and enable amperometric recordings of oxidizable neurotransmitters. The CCEs exhibit excellent electrochemical performance and reproducibility, supporting diffusion-controlled dopamine redox behavior without interference from culture medium. Using ventral midbrain (vMB) organoids at two maturation stages (Day 30 and Day 90), we demonstrate real-time detection of potassium-evoked dopamine release directly from organoids. Mature organoids exhibit higher peak currents, correlating with increased tyrosine hydroxylase (TH⁺) expression confirmed by immunocytochemistry. Embedding organoids in Matrigel significantly attenuates dopamine signals, indicating a diffusion barrier that limits stimulant access and/or dopamine transport to the electrode. Repetitive stimulations reveal progressive depletion of dopamine stores, consistent with diminished vesicular release. Chemical perturbations using 6-hydroxydopamine and verapamil further validate the platform by producing substantial reductions in dopamine release accompanied by loss of TH⁺ neurons. These results establish CCEs as a powerful tool for label-free, real-time monitoring of neurotransmission in human organoids, with potential for multiplexed sensing and integration of enzymatic detection strategies for non-electroactive neurotransmitters.

## 1. INTRODUCTION

Dopamine is a key neuromodulator implicated in reward processing, motor control, synaptic plasticity, and the pathophysiology of neuropsychiatric and neurodegenerative disorders. As other neurotransmitters, dopamine signaling is highly dynamic and occurs at sub-second timescales, underlying the need for analytical tools capable of capturing its release and clearance in real time with both high temporal resolution and chemical specificity [1]

A variety of methods have been developed to quantify dopamine and other neurotransmitters in biological systems, each with important advantages and limitations [2], [3], [4], [5], [6]. Classical techniques, such as high-performance liquid chromatography (HPLC) and mass spectrometry (MS) provide excellent sensitivity and quantitative accuracy. However, require extensive sample handling, e.g., coupling to microdialysis [7], and lack the temporal resolution necessary to resolve rapid release and reuptake events in tissue

Microelectrode arrays (MEAs) are now widely used for real-time electrophysiology in organoids and brain tissue, providing high spatiotemporal resolution of neuronal activity [8], [9], [10]. However, MEAs record changes in extracellular potential rather than the chemical identity of released neurotransmitters, rendering it challenging to distinguish which neurotransmitter triggers a given electrical event.

Optical detection techniques, such as genetically encoded fluorescent reporters or small-molecule probes (so-called sniffer cells), are currently revolutionizing the field, enabling sub-second neurotransmitter measurements in vitro and in vivo [3], [11], [12]. These approaches also face challenges, such as dependence on genetic manipulation of cells or dye loading, restricted optical access, requirement for careful calibration, and sometimes limitations in chemical specificity and photostability in complex tissues.

Electrochemical methods, such as fast-scan cyclic voltammetry (FSCV) and amperometric constant potential recordings, provide high temporal resolution and label-free detection of oxidizable catecholamines like dopamine [5], [13], [14], [15], [16]. However, when employing these methods, one must be aware of susceptibility to background currents, signal drift, electrode fouling, and limited chemical selectivity without using additional molecular recognition elements [13], [17], [18].

Three-dimensional (3D) human brain organoids, including ventral midbrain (vMB) organoids comprising functional dopaminergic neurons, offer physiologically relevant human-derived platforms to study neurotransmission in physiological and pathological context, such as Parkinson’s disease (PD) [19], [20]. However, their size, optical density, and 3D microenvironment pose unique challenges for in situ neurotransmitter measurements. Despite major advances across biochemical, optical, and electrochemical platforms, there is to date no widely adopted technology that combines chemical specificity with real-time sub-second to second detection of dopamine release from intact brain organoids.

We have previously demonstrated real time detection of dopamine release from 2D cultures of human neural stem cell-derived dopaminergic neurons directly on photolithography-based pyrolytic carbon micropillar electrodes and optical fibre electrodes [21], [22]. The height of these micropillar electrodes was around 20 µm, which was sufficient to interact with 2D cell cultures but is not suitable for 3D organoids. In a recent study, we have shown real time recordings of dopamine from vMB organoids using optical fibre electrodes placed in their proximity [16]. However, precise placement of optical fibres in cell culture platforms with multiple organoids is challenging and a more integrated approach is desirable.

This study leverages 3D printing combined with pyrolysis to develop on chip 3D carbon pillar electrodes with heights of several 100 µm [23], and their application for real-time dopamine detection from vMB organoids. A carbon cup electrode (CCE) with a cup-shaped geometry, formed by a multi-level pillar array, is introduced to increase the electrode surface area and to confine a single vMB organoid inside the cup during exocytosis measurements. Real time detection of dopamine in response to chemical stimulation at two distinct developmental stages (Day 30 and Day 90) was demonstrated. The CCE platform enables for the first time capture and detection of a larger portion of dopamine and other oxidizable neurotransmitters in situ and in real-time from organoids, while also paving the way for integrating enzymatic biosensors to detect non-electroactive transmitters such as glutamate and gamma-aminobutyric acid (GABA) [17].

Furthermore, we investigate the impact of embedding organoids in the extracellular matrix (ECM)-substitute Matrigel, commonly used for organoid embedding, revealing that this coating seems to act as a barrier, hampering neurotransmitter detection at the electrode. Additionally, the vMB organoids are subjected to two disruptive treatments (6-OHDA and Verapamil) designed to disrupt exocytotic function [24]. Immunocytochemical analysis was employed to confirm the differentiation of organoids into dopaminergic phenotype.

## 2. RESULTS AND DISCUSSION

### 2.1 From design to reality: Fabricating carbon cup electrodes

The overall process flow of the CCE fabrication, from computer-aided design (CAD) design to the final pyrolytic 3D carbon structure, are detailed in Figures 1A-C. The polymeric precursor structure was 3D printed on an oxidized black silicon substrate (Figure 1B), which had randomly distributed nanograss of about 500 nm in height. This nanostructuring was required to provide adhesion of the CCE to the carrier substrate [23]. The CCE precursor structure was designed with an array of cylindrical pillars, each with a diameter of 100 µm arranged in a circular pattern with 2 mm in diameter (Figure 1D and 1E). The pillar height varied from 650 µm for the inner to 1550 µm for the outer pillars. The pillars were placed on a 50 µm thick base also serving as collector layer and interface for electrode connection (Figure 1F). The simultaneous 3D printing of five electrode precursor structures using the generated CAD was done by projection micro-stereolithography (PµSL) in a chip array mode (Supporting Information, Figure S1A). Figure 1G shows the final pillar cup design, Figure 1H a single 3D-printed cup-shaped precursor structure, Figure 1I the corresponding pyrolyzed CCE, retaining its the original geometry, and Figure 1J and 1K a zoomed-in view of an individual carbon pillar in the CCE.

**Figure 1.**
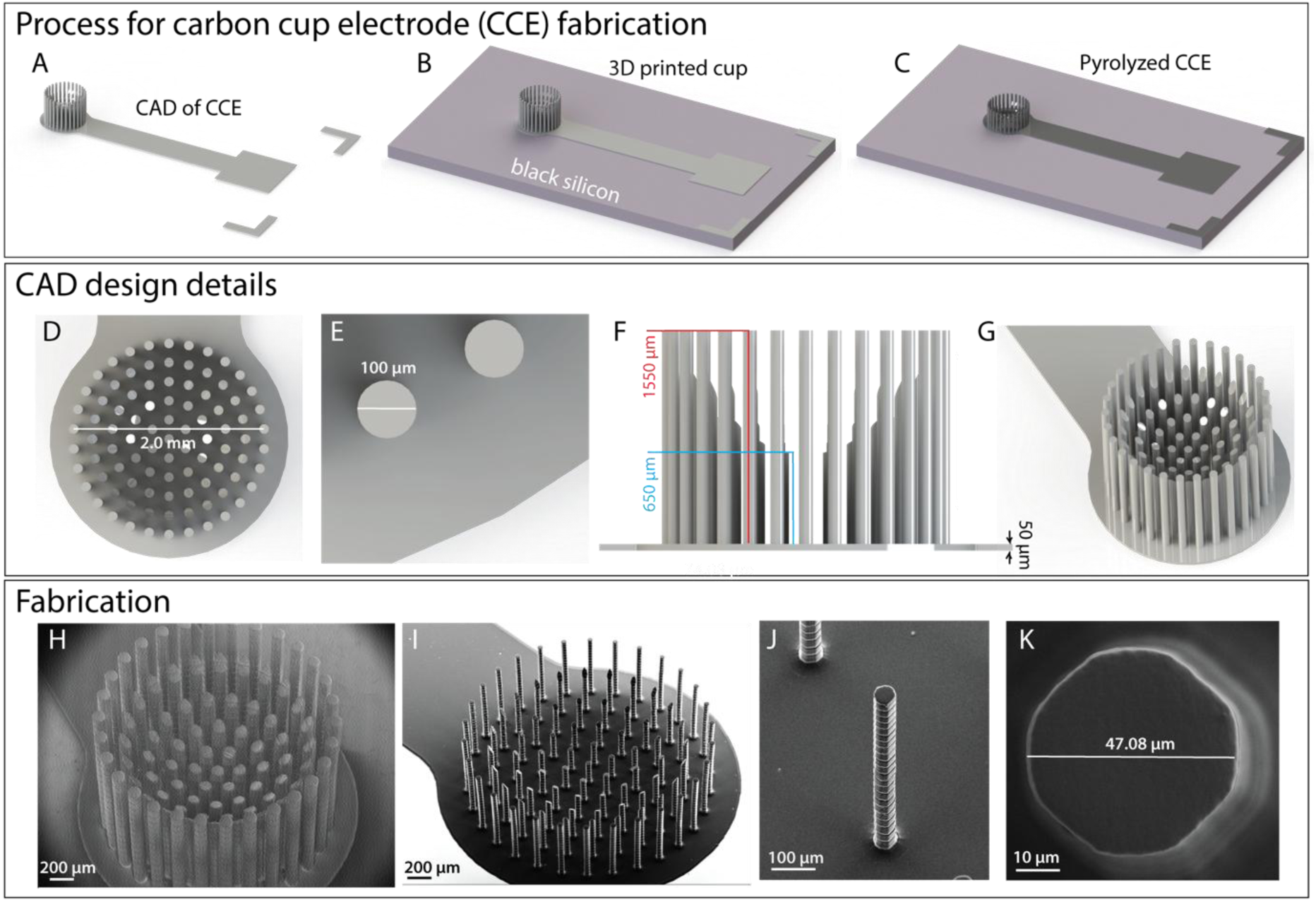
Design and fabrication of CCE. *Top panel:* Stepwise fabrication of a CCE on a black silicon substrate: A) CAD design of CCE, B) 3D printed precursor cup and C) pyrolyzed CCE. *Middle panel:* CAD design details: D) Top view of cylindrically distributed pillars and E) zoom in on two single pillars. F) Side view, featuring pillars with variable heights ranging from 650 to 1550 µm. G) Final cup shaped structure. *Bottom panel:* SEM images of fabricated structures: H) 3D printed polymer precursor cup. I) Final pyrolyzed CCE with zoom in on a single carbon pillar, side view (J) and top view (K).

During pyrolysis, the organic structure of the polymeric precursor is disrupted resulting in mass loss and increase in the carbon density [25]. The thermal profile of the pyrolysis process included three initial steps at temperatures of 340 °C, 400 °C and 425 °C, respectively, before final ramping to 900 °C. This stepwise pyrolysis is crucial since most of the resin is degraded between 350-450 °C and a controlled mass loss is imperative to prevent structural deformation of the carbon electrodes [26]. As an inherent feature of the pyrolysis process, shrinkage of the CCE compared to the cup-shaped 3D printed structures, both in pillar height (60%) and diameter (50%), was observed. However, the flat base structure did not shrink laterally because it was restricted by the black silicon substrate, resulting in a 95% reduction in height only to a carbon thickness of 2.5 µm.

### 2.2 A closer look: Dopamine detection using carbon cup electrodes

The electrical properties of pyrolytic carbon electrodes depend on both the polymer precursor, the 3D printed geometry and the pyrolysis conditions used during fabrication. The manufacturing process showed excellent reproducibility, as evidenced by highly consistent cyclic voltammograms (CVs) obtained from different CCEs within and across chip arrays (Figure S2). Prior to all electrochemical measurements, the electrodes were treated with oxygen plasma to enhance performance. This treatment introduces oxygen-containing functional groups that promote electron transfer and increase the hydrophilicity of the carbon surface [27].

CVs for 150 µM dopamine (Figure 2A) confirm a quasi-reversible redox reaction in accordance with other studies using different polymer precursors for carbon electrode fabrication [28]. A linear dependence of the peak current with the square root of the scan rate (Figure 2B) indicates a diffusion-controlled process. Moreover, the linearity even at higher scan rates suggests that the electrochemical performance is not limited by the resistivity of the pyrolytic carbon derived from the 3D printed precursor structures.

**Figure 2.**
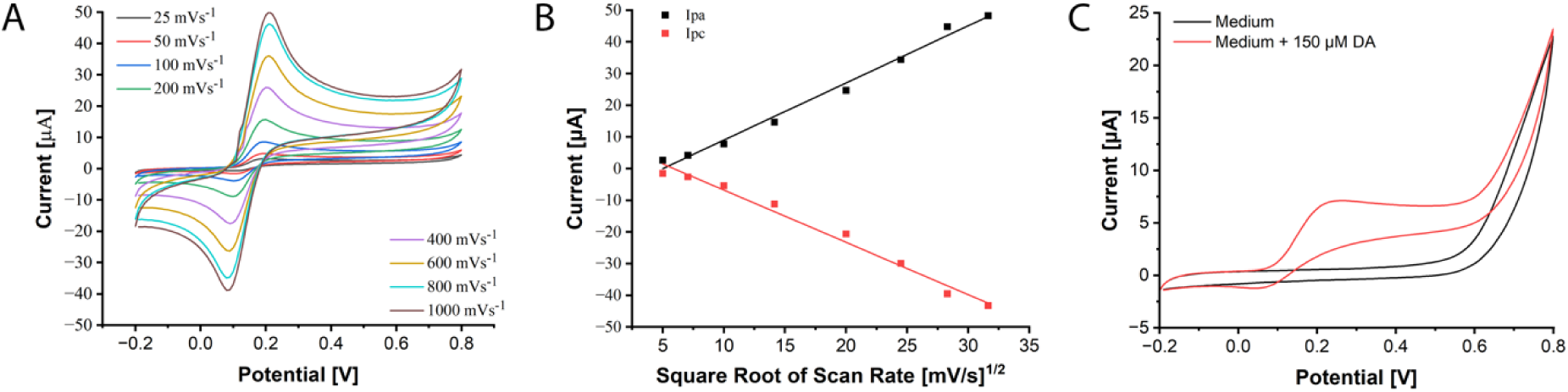
A) CVs at different scan rates recorded for CCEs vs Aga/AgCl for dopamine (150 µM) in PBS (pH 7). B) Peak current versus square root of scan rate from anodic and cathodic current peaks. C) CVs recorded at a scan rate of 100 mV s-1 in Neurobasal medium with and without added dopamine (150 µM).

In many studies, depolarization of neurons is typically induced by elevated K^+^ concentration in a buffer solution [29]. In our study, Neurobasal medium (Gibco) was used to avoid medium exchange during measurement and to better mimic the cell culture environment during baseline recording and exocytosis. Experiments were carried out to investigate any potential interferences caused by components of the cell culture medium used for the vMB organoids. CVs acquired in Neurobasal medium (Figure 2C) show no presence of faradaic processes in the potential range relevant for dopamine detection, whereas the same medium spiked with dopamine shows the presence of a small anodic peak. Comparative CVs acquired in neurobasal medium and PBS (Figure S3) show a marginally higher capacitive current for neurobasal medium in the applied potential range and an elevated faradaic current at potentials above 0.6 V (vs. RE), which are beyond the potential applied for dopamine detection (0.5 V). This absence of interferences suggests that the Neurobasal medium is suitable for stimulation of neurons in organoids without influencing dopamine detection.

### 2.3 Developing vMB organoids and assessing their dopaminergic profile

vMB organoids were generated from human embryonic stem cells (hESCs, RC17 line) cultured in 96-well ultra-low attachment plates following the procedure developed by Corsi et al [30] illustrated in Figure 3A. After 2 days of culture, cell aggregates were observed and the cell culture medium was replaced with the differentiation medium. On day 16, developing vMB organoids (embryoid bodies) were separated into two groups: one group was embedded in Matrigel and the other was kept non-embedded. Both embedded and non-embedded organoids were transferred to ultra-low attachment 6-well plates until organoids were either 30 or 90 days old.

**Figure 3.**
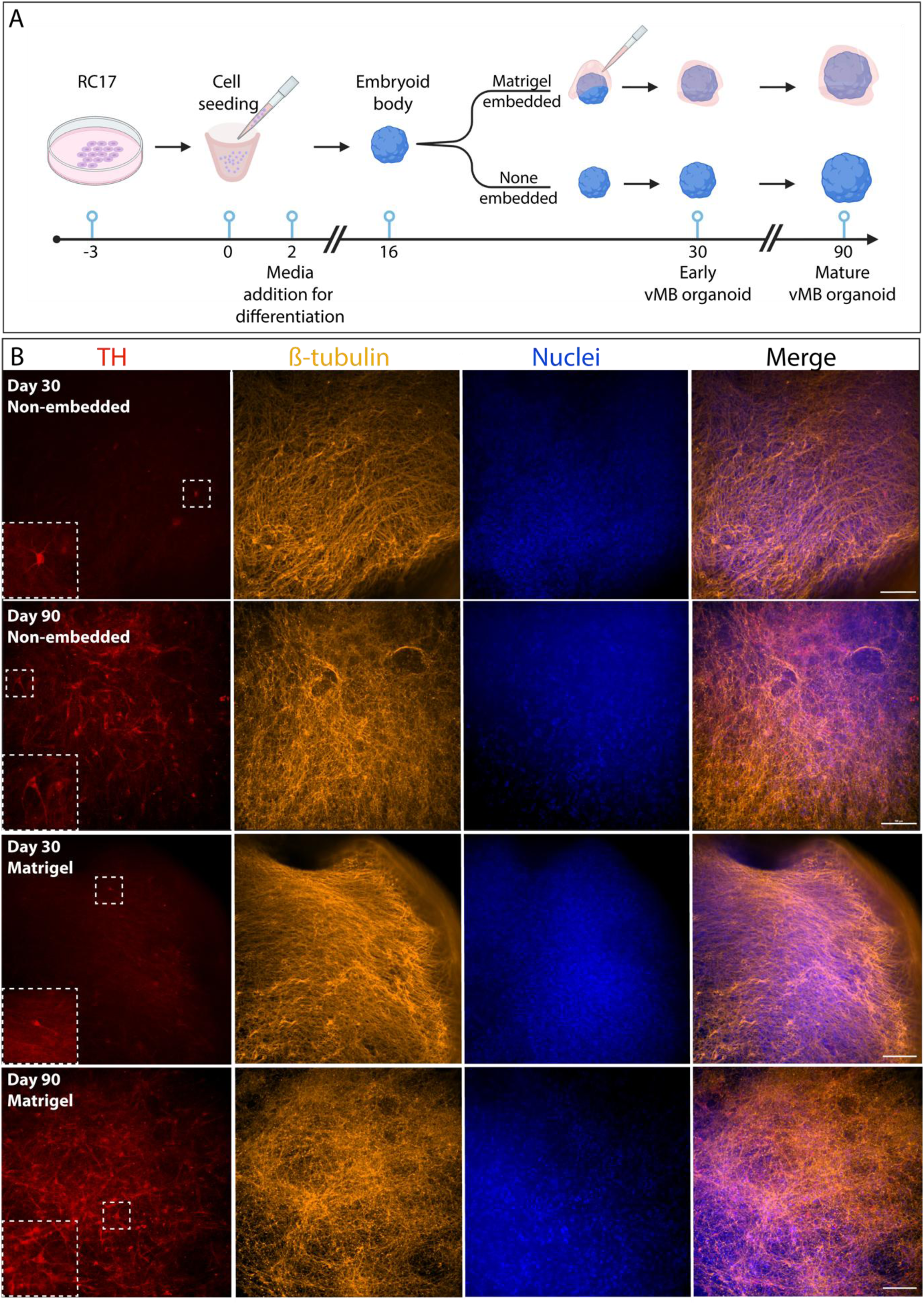
**A)** Procedure for generating vMB organoid cultured until day 30 and 90 with and without Matrigel embedding initiated at day 16. B) Immunocytochemistry of non-embedded and Matrigel embedded vMB organoid at day 30 and 90. A few TH⁺ cells are present at day 30, indicating early maturation, which by day 90, has markedly increased for both cases, indicating more mature vMB organoids at day 90. Large white dotted box: enlarged image of TH positive cells in area of smaller, dotted box. Scale bar = 100 µm.

Dopaminergic (DA) neurons are the predominant neuronal population in vMB organoids [16], and immunocytochemistry (ICC) was performed to confirm their presence (tyrosine hydroxylase positive (TH⁺) cells) at the two time points, for both non-embedded and Matrigel embedded organoids (Figure 3B). The representative ICCs show just a few of TH⁺ cells at day 30, consistent with early stages of maturation. By day 90, the number of TH⁺ cells have substantially increased, indicating that both non-embedded and Matrigel embedded vMB organoids have reached maturation.

### 2.4 Capturing real-time dopamine release from vMB organoids in a CCE

Amperometric real-time detection of dopamine release from silk scaffolded vMB organoids was previously demonstrated using a carbon-coated fiber electrode [16]. In this study, we recorded dopamine exocytosis at day 30 and day 90 for embedded and non-embedded vMB organoids with a single organoid carefully pipetted into a CCE (Figure 4A and B, Video S1). A custom-made holder with a three-electrode setup was designed to accommodate one CCE array chip (five CCEs as working electrodes (WEs)), each with a vial for cell medium, space to position an external counter (CE) and reference electrode (RE), as well as a printed circuit board (PCB) containing spring loaded pins to interface the CCE array chips (Figure 4C, Figures S1A and S1B).

**Figure 4.**
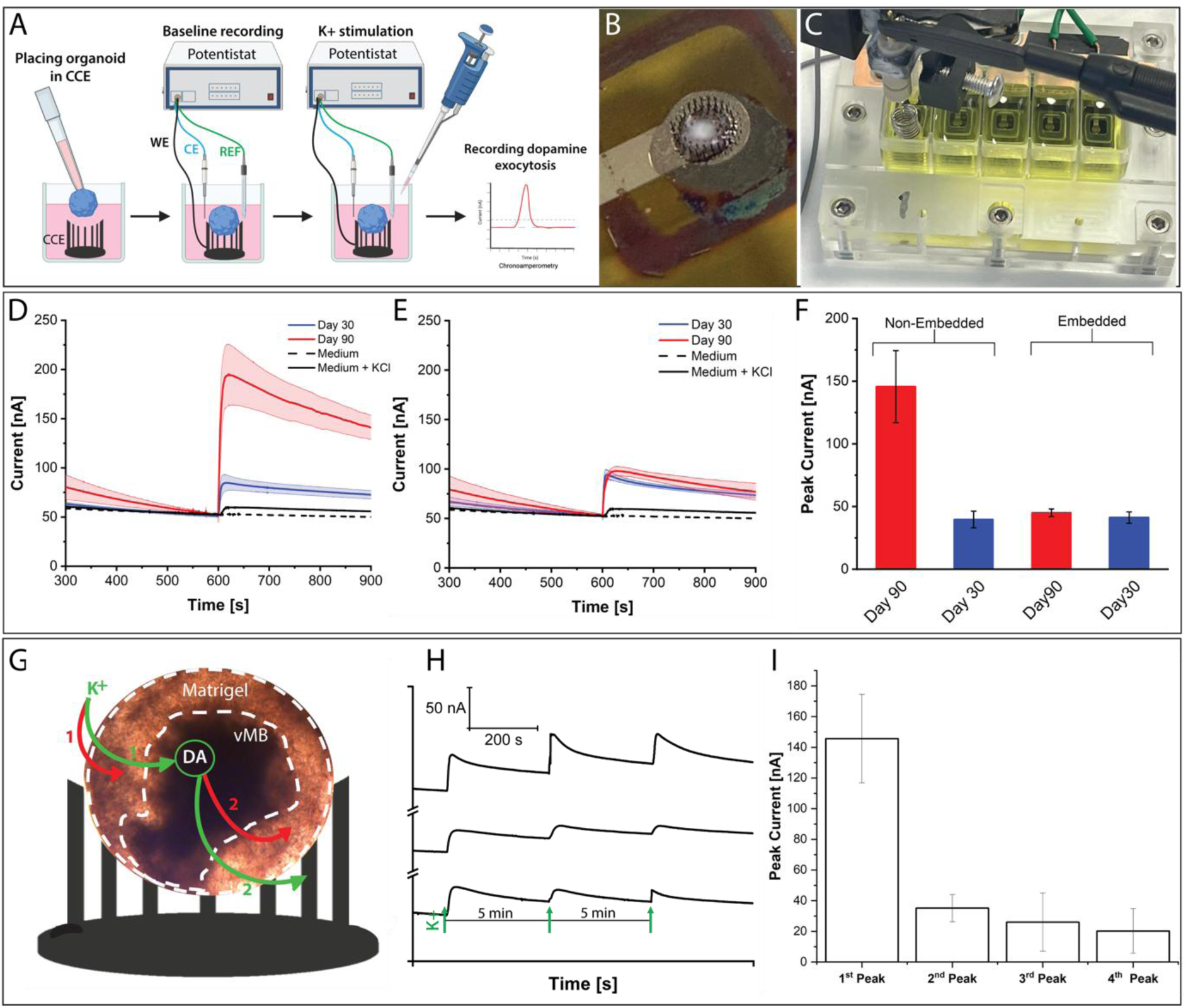
*Top panel:* A) Procedure for amperometric dopamine exocytosis measurements of organoids placed in CCEs. B) Photo of a vMB organoid placed in a CCE. C) Custom made measurement set up for an array chip housing five CCEs. *Middle panel:* Amperometric recordings of potassium evoked dopamine release from vMB organoids (n=3) at Day 30 (blue) and Day 90 (red) of maturation: D) non-embedded (n=3), E) Matrigel embedded vMB organoids, and F) bar plots comparing peak currents for embedded and non-embedded organoids at different days. *Bottom panel:* G) Photograph of a Matrigel embedded vMB organoid placed in a schematic CCE: white dotted lines show Matrigel embedding the organoid, stimulation with K^+^ and its path through the barriers, reaching the vMB (green arrow) or not (red arrow), stimulating release of dopamine (DA) and its path through the barriers (green arrow) or not (red arrow) for detection by the CCE (green arrow). H) Repetitive potassium stimulation of three different non-embedded vMB organoids (day 90) at an interval of 5 minutes during continued amperometric recordings of the resulting dopamine release, green arrows indicating time for K^+^ stimulation. I) Bar plots of the peak height decrease with respect to the baseline after each stimulation. The 1^st^ column shows the average peak height of the first stimulation shown in Figure D (red) performed 2 hours prior the subsequent repetitive stimulations.Error bands in Figures D and E represent standard deviation of measurements (n=3).

After recording the baseline, dopamine exocytosis was initiated by adding Neurobasal medium with elevated K^+^ concentration, while continuing the amperometric recording for 20 min. Figure 4D demonstrates successful in situ detection of dopamine release from non-embedded vMB organoids. The recordings show that the dopamine peaks at both Day 30 and Day 90 rise quickly but drop very slowly back to the baseline. This prolonged dopamine release from the organoids, compared with measurements on 2D cell populations [21], is due to the complex nature of organoids. Dopaminergic neurons are scattered throughout the 3D matrix and require a longer period for the stimulation medium to reach the cells that are not on the surface of the organoid, and equally for dopamine to diffuse out to be oxidized at the CCE. The non-embedded vMB organoids show clear and distinct peaks for both timepoints with the highest response at day 90 (Figures 4D and 4F). This is expected since dopaminergic neurons should not be fully differentiated at day 30.

The influence of Matrigel on vMB organoids has not been thoroughly investigated, except for its influence on early-stage development of cerebral and telencephalic organoids [31], [32]. We investigated if Matrigel embedding had any influence of recording dopamine exocytosis from vMB organoids. After recording the baseline, dopamine exocytosis was initiated by adding Neurobasal medium with elevated K^+^ concentration, while continuing the amperometric recording for 20 min. Figures 4D and 4E show that dopamine current peaks for both embedded and non-embedded organoids rise quickly and drop very slowly back to the baseline. However, embedded organoids show significantly lower and overlapping peaks for Day 30 and 90 (Figures 4E and 4F) compared to those cultured without Matrigel, reaching similar peak heights as for non-embedded organoids at day 30. This suggests that Matrigel hinders dopamine detection. One scenario is that the K^+^ stimulation medium does not diffuse through the Matrigel to reach the cells inside the organoid; hence, they cannot be depolarized (Figure 4G, red arrow 1). Alternatively, the cells are depolarized (Figure 4G, green arrow 1) but the released dopamine is captured inside the Matrigel layer and thus prevented from reaching the CCE (Figure 4G, red arrow 2). In either scenario, Matrigel seems to act as a barrier between the cells and the electrode, impeding the electrochemical response that is strongly dependent on facile mass transfer of dopamine to the electrode surface (Figure 5G, green arrow 2).

**Figure 5.**
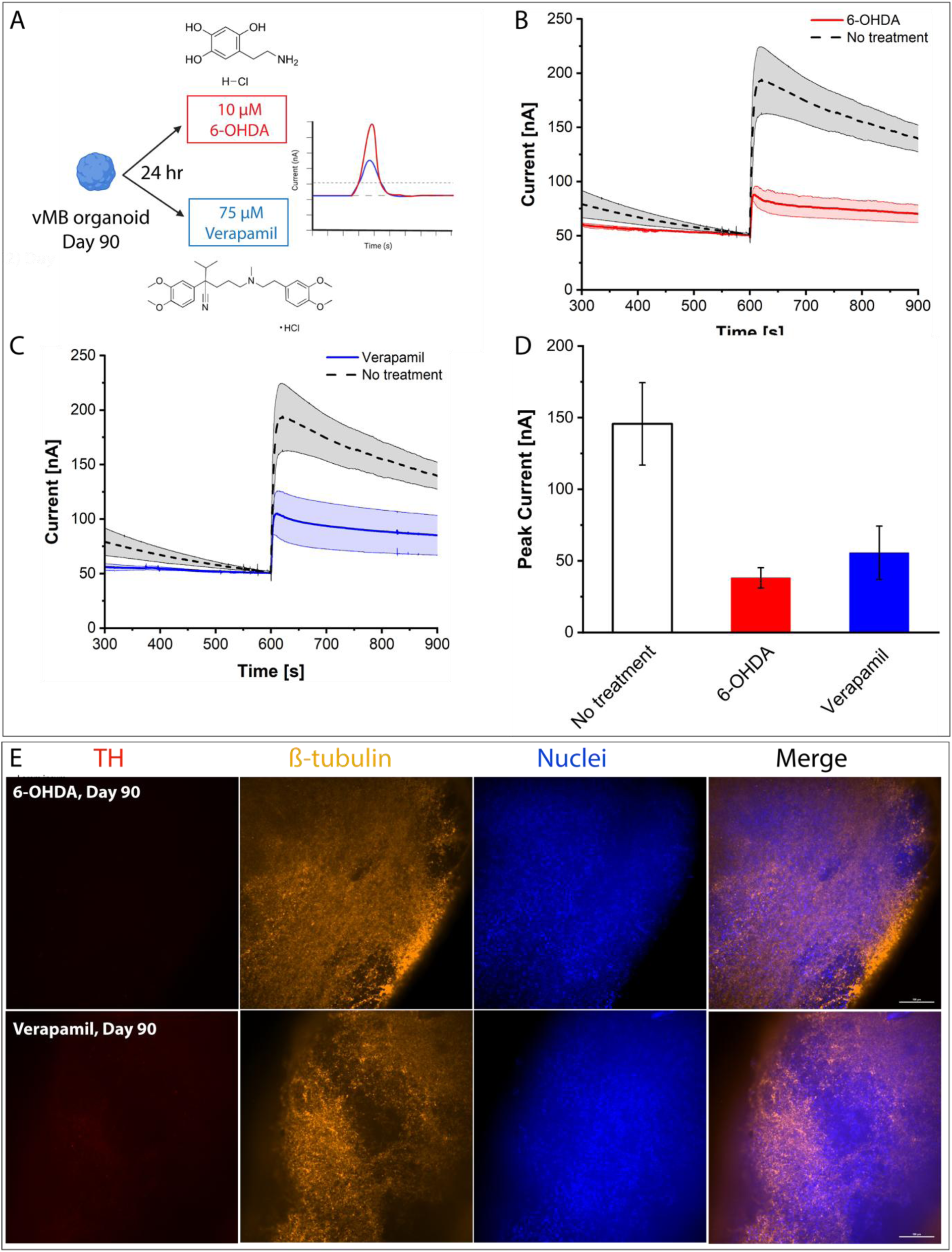
A) Dopamine exocytosis measurements from non-embedded vMB organoids subjected to chemical treatments at day 90 (6-OHDA and Verapamil). Amperometric recording before and after treatment for three different vMB organoids: B) 6-OHDA, C) Verapamil, and D) bar plots comparing peak currents for both treatments (n=3). E) Immunocytochemistry of vMB organoid after treatment with 6-OHDA and Verapamil at day 90. TH (red), β-tubulin (orange), nuclei (blue), and merged images (red, orange, blue). Scale bar = 100 µm. Error bands in Figures B and C represent standard deviation of measurements (n=3).

To investigate the sustainability of exocytosis, an additional experiment was conducted: Dopamine release was repetitively triggered in non-embedded organoids at Day 90. Exocytosis of three separate activations were recorded for three different organoids (Figure 4H and 4I). It should be noted that the organoids placed in the CCEs used for this experiment were the same as those previously stimulated with increased K^+^ concentration (1^st^ stimulation shown in Figure 4D and 4I) but with a resting period of 2 hours before the experiment. Therefore, peaks 2-4 observed in Figures 4H and 4I have a lower current amplitude than what is observed in the measurements shown in Figure 4D and 4I. For each subsequent activation, the dopamine release diminishes due to vesicle pool depletion (Figures 4H and 4I), similar as to what was previously observed using optogenetically modified ventral mesencephalic stem cells (hVM1) differentiated on a carbon-coated optical fiber electrode [22].

### 2.5 Impact of neurofunctional modulators on dopamine release from vMB organoids

To investigate how known neurofunctional modulators affect the exocytotic machinery of dopamine, two pharmacological treatments were applied to non-embedded vMB organoids at Day 90 (Figure 5A). The first treatment involved 6-OHDA, which is well-known to induce neurodegeneration and death of TH^+^ neurons through oxidative stress [33]. The amperometric peak heights for 6-OHDA treated organoids show a distinct decrease (Figure 5B), reaching a similar level as the baseline current for untreated organoids. The second treatment was performed using Verapamil, which acts as a blocker of voltage-gated calcium channels, preventing calcium influx into depolarized cells [34]. Verapamil-induced blocking of calcium influx leads to inability to initiate fusion of neurotransmitter vesicles with the plasma membrane, which is required for exocytotic neurotransmitter release. Moreover, prolonged exposure to verapamil has been shown to disrupt intracellular calcium homeostasis, ultimately leading to apoptosis and death of neurons [35]. The amperometric peak height decreased significantly compared to untreated organoids (Figure 5C). Both chemical treatments resulted in decreased dopamine release (Figure 5D) verifying that the DA release recorded is DA neuron and activity specific; however, the precise mechanism of verapamil warrants further investigation. ICC of organoids having undergone these two treatments clearly corroborates these findings (Figure 5E). None or only very few TH⁺ cells are observed after treatment with chemicals compared with the untreated condition (Figure 3B, Day 90).

## Conclusions

This study showcases the use of carbon cup electrodes (CCE), derived from 3D printing, for real-time monitoring of dopamine release from vMB organoids. A cup-shaped design, featuring multi-level cylindrical pillars, was implemented to confine a single vMB organoid inside the cup, increase the electrode surface area, and capture dopamine in a 3D configuration from a larger region of the organoid. Additionally, the cup design prevents organoid movement during injection of the stimulation buffer. The CCE demonstrated an outstanding electrochemical performance. TH expression, confirming the presence of dopaminergic neurons in vMB organoids at different maturation stages, was demonstrated using immunocytochemistry. The vMB organoids were generated with and without embedding in an ECM layer of Matrigel to investigate if the embedding had any influence on dopamine detection. Amperometric recordings of dopamine exocytosis from potassium stimulated vMB organoids showed significant decrease in current when the organoids were embedded in Matrigel, indicating that Matrigel introduces a diffusional barrier, hindering the access of dopamine to the CCE. Organoids were also subjected to disruptive pharmacological treatment with 6-OHDA and Verapamil, showing the expected reduction of dopamine release accompanied by significantly lower peak currents obtained than for untreated organoids at the same maturation level. The results highlighted in this paper show the ability of the CCE to serve as a platform for real-time dopamine detection from vMB organoids. Future possibilities include chemical modifications to introduce biosensor capabilities for detection of non-oxidizable neurotransmitters to study different region-specific organoids, as well as fabrication of CCEs incorporating individually addressable carbon pillar electrodes for multiplexed neurotransmitter detection in individual organoids or assembloids used for disease modelling applications.

## 3. MATERIALS AND METHODS

### 3.1 Fabrication of carbon cup electrodes (CCE) and measurement system

#### Substrate preparation

A nanograss surface, also known as black silicon, was created on a 150 mm silicon wafer (Siegert Wafers, Germany) using deep reactive ion etching (DRIE, Pegasus, SPTS Technologies Ltd., Newport, UK). A combination of oxygen and sulfur hexafluoride gases based on the CORE process [36] was used. The black silicon wafer was then oxidized using a tube furnace (Tempress, The Netherlands) in oxygen atmosphere at 1050 °C for 2 h, resulting in 130 nm of thermally grown silicon oxide serving as insulating layer. The wafer was finally cut in 54.0 x 19.2 mm pieces using a dicing saw (DAD 3421, Disco, Japan).

#### *CAD design of CCE electrode* (Figure 1A-G)

The 3D model was designed with a computer aided design (CAD) software (SolidWorks, Dassault Systèmes, France) and converted into 2D images, each image comprising a 5 µm layer of the model using Voxeldance Additive software (Shanghai, China).

#### Polymer precursor structure printing

The black silicon substrate was fixed on a printing platform using double adhesive tape (AR7840, Adhesive Research, USA). A projection micro-stereolithography (PµSLA) printer (microArch S240, BMF, USA) was used to print five electrode precursor structures in the chip array mode, using a CAD design (Figure S1A) and a high temperature resin (HTL-20, BMF, USA) similarly as reported earlier [23]. The 3D structures were printed with a layer thickness of 5 µm layers. The light intensity and exposure time were optimized according to the layer thickness (Table S1). The total printing time for five cup-shaped structures was 12 h. The electrode precursor chip array was thoroughly washed with isopropanol (IPA) for 5 min, and then carefully released from the platform using a scalpel followed by backside cleaning with a wipe and acetone.

#### *Pyrolysis* (Figure 1J-K)

Three chip arrays each containing five precursors cup structures were placed on a silicon carrier spaced 10 mm apart inside a quartz tube furnace (MTI Corp., USA) with a diameter of 60 mm. The silicon carrier was placed such that the central chip was positioned in the middle of the tube (Figure S2A). The thermal process was carried out with a constant flow of 400 sccm nitrogen gas. The temperature was ramped in a stepwise manner previously optimized using thermal gravimetric analysis (TGA) [26]. The initial ramping step started from room temperature and increased to 340 °C at 2 °C min^-1^ where it was held for 1 h. In the second step the temperature was increased to 400 °C at 2 °C min^-1^ and held there for another hour. In the third step, the temperature was increased to 425 °C at the same heating rate and holding time. The final step involves a ramp up process - the graphitization step - where the temperature was raised to 900 °C at 2 °C min^-1^where it was held for 1 h. A cooling down step was performed by turning off the heaters and waiting for the furnace to reach 50 °C before unloading the samples. The entire pyrolysis process, from the initial ramping up to cool down, took about 12 h. To evaluate the shrinkage during pyrolysis, both the precursor cup structures and CCE were imaged using a scanning electron microscope (SEM; Supra 40vp (Zeiss, Germany) using high vacuum mode (variable pressure mode (VPSE mode)) at 25 Pa.

#### Electrode holder and PCB

A polymethylmethacrylate (PMMA) based custom-made holder was designed and micro-milled to accommodate one CCE array chip, each with a vial for cell medium, space to position an external CE and RE electrode, as well as a PCB containing spring loaded pins to interface the electrodes (Figures 4C and S1B). The five CCEs within the chip array were individually separated by the holder in a manner that each CCE was electrically and fluidically insulated from each other. The holders were designed in CAD software and their fabrication outsourced to JLCPB (China). PMMA was chosen due to its biocompatibility and chemical resistance to the sterilization process. To ensure fluidic sealing, 1 mm thick silicone O-rings (RCT Reichelt Chemietechnik GmbH, Germany) were laser cut using an Epilog Mini 18 CO_2_ laser system (Epilog Laser, Golden, CO, US).

### 3.2 Characterization of carbon cup electrodes (CCEs)

The CCEs were oxygen plasma treated for 75 s at a pressure of 0.6 mbar and power of 120 W, using a Zepto plasma system (Diener GmbH, Germany).

A three-electrode setup was employed with the CCE as WE situated at the bottom of the vial (Figure S1B), and a 500 µm diameter platinum wire (Advent Research materials Ltd., Oxford, England) and a DRIREF-L Ag|AgCl (saturated KCl) (WPI, Sarasoata, FL, US) as external CE and RE, respectively, inserted into the vials from the top (Figure 4C). Electrode characterization was performed using cyclic voltammetry in 150 µM dopamine solution prepared in de-aerated PBS (pH 7.0) (Figure 2). A PalmSens4 potentiostat (PalmSens BV, Netherlands) was used to sweep the potential from -0.2 to 0.8 V at different scan rates (from 25 mV s^-1^ to 1 V s^-1^). CVs were acquired in Neurobasal medium with and without added dopamine at a fixed scan rate of 100 mV s^-1^. To evaluate if the position of three precursor array chips, when placed simultaneously in the furnace (Figure S2A), influences the electrochemical proprieties of the resulting individual arrays of 5 CCEs, CVs from all 15 CCEs were acquired in 10 mM hexacyanoferrate (II/III) ([Fe(CN)_6_]^-4/-3^) in PBS and compared to CCEs in a chip array pyrolyzed individually (Figure S2B). Peak analysis was carried out by extracting current peak heights (i_pa_, i_pc_) and peak-to-peak potential separation (ΔE_p_).

### 3.3 Developing vMB organoids and immunocytochemistry analysis

*vMB organoids* (Figure 3A): RC17 cells (Roslin Cells, hESCs, cat. no. hPSCreg RCe021-A) were seeded on a 6-well tissue culture plate pre-coated with growth factor reduced (GFR) Matrigel (cat. no. 356230) at a concentration of 0.5 µg in iPS brew medium (Miltenyi, cat. no. 130-104-368). When cells reached a confluency of 70%, they were passaged. Cells were then cultured in iPS-brew medium with Y-27632 supplement for 24 h. For cell culture maintenance, the medium was replaced with fresh iPS-brew every day. vMB organoids were generated according to Sozzi et al [37]. RC17 cells, maintained in a 6-well plate, were treated with 50 µl cm^-2^ Accutase (Thermo Fisher Scientific, cat.no. A1110501) for 5 min at 37°C. 1.5 mL of washing medium, containing 5% knock-out serum replacement (KOSR) (Thermo Fisher Scientific, cat.no. 10828010) in DMEM/F12 (Thermo Fisher Scientific, cat.no. 10565018) was then gently added (without removing Accutase) to detach and harvest the cells. A single cell suspension of 8,000 cells/well in 200 µL iPS-Brew per well, supplemented with 10 µM Y-27632 (Tocris Bioscience, cat. no. 1254) was pipetted into an ultra-low attachment 96-well plate (Corning Costar, cat. no. 7007). The plate was left undisturbed in an incubator for 48 h, after which the medium was replaced with fresh iPS-Brew without Y-27632. After 2 days of culture the medium was replaced with differentiation medium, containing 1:1 DMEM/F12: Neurobasal medium (Thermo Fisher Scientific, cat.no. 10565018 and cat.no. 10888022), 1:100 N_2_ supplement (Thermo Fisher Scientific, cat.no. 17502048), 10 µM SB431542 (Tocris Bioscience, cat. no. 1614), 150 ng mL^-1^ rhNoggin (Miltenyi, cat. no. 130-103-456), 400 ng mL^-1^ SHH-C24II (Miltenyi, cat. no. 130-095-727), 1.5 µM CHIR99021 (Tocris Bioscience, cat. no. 4423), 1:100 MEM Non-essential amino acids solution (Thermo Fisher Scientific, cat.no. 11140050), 1:1000 2-Mercaptoethanol (Thermo Fisher Scientific, cat.no.31350010), 1:500 Penicillin-Streptomycin (Thermo Fisher Scientific, cat.no. 15140122), which was maintained for the entire differentiation period with medium exchange on day 3 and day 7. On day 8, the medium was replaced with the previous medium supplemented with fresh 100 ng mL^-1^ FGF8b (Peprotech, cat no. 100-25-25UG). On day 11, this medium was replaced with Neurobasal medium supplemented with 1:50 B27 without vitamin A (B27 medium) (Thermo Fisher Scientific, cat.no. 12587010), fresh 100 ng mL^-1^ FGF8b, 20 ng mL^-1^ BDNF (Peprotech, cat no. 450-02-10UG), and 200 µM ascorbic acid (Sigma Aldrich, cat no. A4403). To embed organoids in Matrigel, they were transferred from the ultra-low attachment plate to a dimples parafilm, using a cut P200 tip (ThermoFisher Scientific), with subsequent removal of excess medium. The organoids were then covered with 30 µL Matrigel and incubated for 30 min to allow crosslinking. Both embedded and non-embedded organoids were transferred to ultra-low attachment 6-well plates each plate containing approximately 10 organoids with terminal differentiation in Neurobasal medium containing 1:50 B27 supplement, 1:100 Glutamax (Thermo Fisher Scientific, cat.no. 35050061), fresh 20 ng mL^-1^ BDNF, 10 ng mL^-1^ GDNF (Peprotech, cat no.450-10-10UG), 200 µM ascorbic acid, 500 µM cAMP (Sigma-Aldrich, cat. no. D0627), and 1 µM DAPT (Tocris Bioscience, cat. no. 2634). During terminal differentiation, 4 mL of medium per well were replaced with fresh medium 2 to 3 times per week.

#### Immunocytochemistry analysis

Both embedded and non-embedded vMB organoids were washed 3 times for 10 min with DPBS without calcium or magnesium (DPBS-/-) (Thermo Fisher Scientific, cat.no. 14190144). Organoids were then fixed with 4% paraformaldehyde (Sigma-Aldrich, cat no. 1.00496) for 1 h at room temperature followed by washing 3 times for 10 min with DPBS-/-. Permeabilization of organoids was achieved overnight using a blocking buffer, containing 5% normal goat serum (ThermoFisher Scientific, cat.no. 31872) and 0.5% triton X-100 (Sigma Aldrich cat no. X100) in DPBS-/-. Primary antibodies, dissolved in blocking buffer, (rabbit polyclonal anti-tyrosine hydroxylase (TH; 1:500; Pel-Freez Biologicals cat.no. P40101), rabbit polyclonal anti-glutamic acid decarboxylase 65/67 (GAD65/67; 1:500; Sigma-Aldrich cat.no. G5163), and mouse monoclonal anti-β-tubulin (TUBB3; 1:500; Sigma-Aldrich cat.no. T8660) were then applied and incubated overnight. The organoids were thereafter washed 3 times for 10 min with DPBS-/- to remove primary antibodies. Secondary antibodies (Alexa 546 goat anti-mouse; 1:1000; ThermoFisher Scientific, cat.no. A11030, Alexa 647 goat anti-rabbit; 1:1,000; ThermoFisher Scientific, cat.no. A21245), and nuclear stain (Hoechst33342; 1:1,000: ThermoFisher Scientific, cat.no. 62249) were then added and incubated overnight. The organoids were finally washed 3 times for 10 min each with DPBS-/-. Images were captured using a spinning disk confocal microscope (Nikon, Japan), and image analysis was conducted using ImageJ software (Figure 3B and 5E).

### 3.4 Real-time detection of dopamine release from vMB organoids in a CCE

Dopamine release from organoids, by K^+^ evoked chemical stimulation, was measured using amperometry with a three-electrode setup (Figure 4A) with a CHI 1010 potentiostat (CH Instruments, Austin, TX, US). A constant potential of 0.5 V (vs. RE) with a current sampling interval of 100 ms was applied and the baseline current was recorded for 10 min in 1.4 mL of fresh Neurobasal medium for terminal differentiation. While continuing the amperometric recording, dopamine exocytosis was then initiated by adding 0.7 mL of the same medium containing 450 mM K^+^ into the vial housing the vMB organoid to elevate the K^+^ concentration to 150 mM. The release of dopamine was detected immediately after stimulation, and the amperometric recording was continued for 20 min (Figure 4D and 4E). For repetitively triggering of dopamine release (Figure 4H and 4I), the same non-embedded organoids at Day 90 were used (Figure 4D, red curve), however first after resting in the incubator for about 2 h. At this point 1/3 of the total medium volume was removed from the vial after each stimulation and replaced with 1/3 of fresh K^+^ Neurobasal stimulation medium to maintain the same conditions as for the previous activation. Each repetitive K^+^ stimulation was performed 5 min apart at room temperature.

### 3.5 Pharmacological treatment of vMB organoids

The vMB organoids were treated with two compromising chemical reagents, 6-OHDA and Verapamil (Figure 5): 6-OHDA was freshly prepared by dissolving 6-OHDA powder (Sigma-Alrich, cat no. H4381) in filtered 0.9% saline solution, containing 0.1% ascorbic acid, to obtain a 10 mM stock solution. The vMB organoids were then treated with 10 µM 6-OHDA for 24 h. Verapamil was prepared by dissolving Verapamil powder (Tocris Bioscience, cat. no. 0654) in sterile water to a stock solution of 5 mM. The vMB organoids were treated with a final Verapamil concentration of 75 µM in medium for 24 h. After 24 h, the treated organoids were placed in the CCEs and exocytosis measurements were performed using the same procedure as described above.

## Acknowledgements

We gratefully acknowledge funding from the HORIZON-EIC-2021-PATHFINDEROPEN-01 project 101047177 OpenMIND and HORIZON-ERC-2024-SYG, project 101167102 Custom-Made. SSK would like to acknowledge financial support by the European Research Council under the Horizon 2020 Framework Programme grant no. 772370-PHOENEEX. The authors acknowledge PolyFabLab, a research infrastructure financially supported by the Novo Nordisk Foundation, grant number NNF21OC0068814.

