## Supporting Information for "Real-Time Neurotransmitter Sensing in Midbrain Organoids Using 3D Carbon Cup Electrodes"

### Supporting Information 1

#### Fabrication and electrochemical characterization of CCE

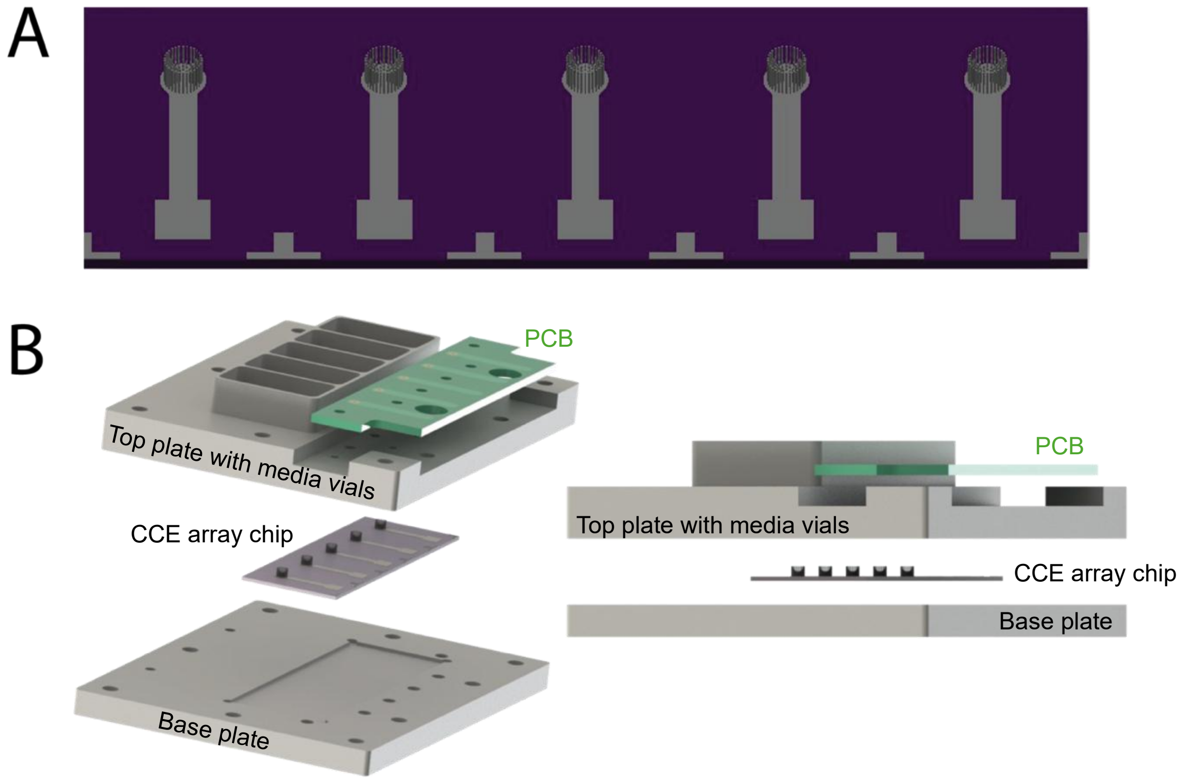


**Figure S1.** A) CAD designs of an CCE array chip with 5 CCEs. B) CAD exploded top and side view of the electrochemical measurement set-up including base plate, top plate with medium vials, and a PCB accommodating one chip array.

**Table S1.** Parameters for 3D printing CCE on chip. All parameters are based on Boston Microfabrication (BMF) S240 printer. Intensity is an integer (max value = 255) for a maximum intensity of 90 $mWcm^{-2}$. The layer thickness is defined by P-d – P-u values. Dwell time 1 and 2 for when the stage moves downwards and upwards, respectively. The first layer has a slightly higher intensity to enhance the adhesion to the substrate. The layer thickness is 5 μm for all layers.

| | **Image** | **Exposure Time (s)** | **Intensity** | **P-d (mm)** | **Dwell Time 1 (s)** | **P-u (mm)** | **Dwell Time 2 (s)** | **Scrapper** |
|---|---|---|---|---|---|---|---|---|
| 1 | 1 | 1 | 40 | 0 | 0 | 0 | 300 | 0 |
| 2 | 10 | 1 | 30 | 4 | 2 | 3.995 | 300 | 2 |
| 3 | 309 | 1 | 30 | 4 | 2 | 3.995 | 60 | 10 |

**A**

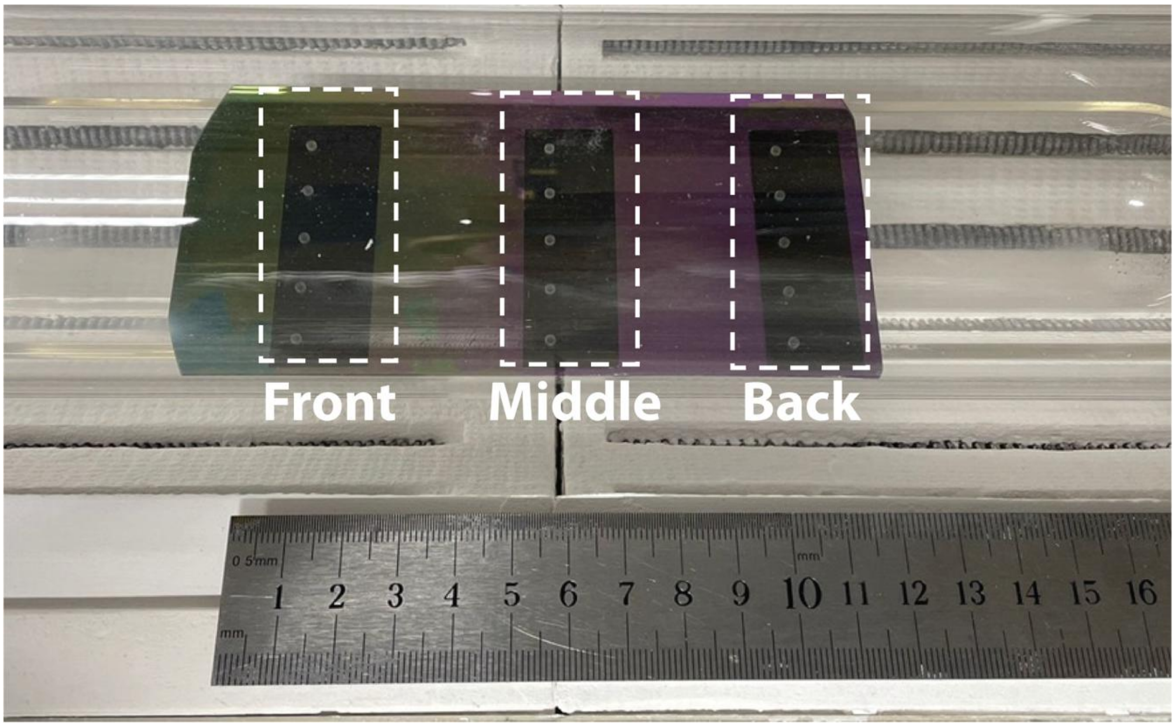


**B**

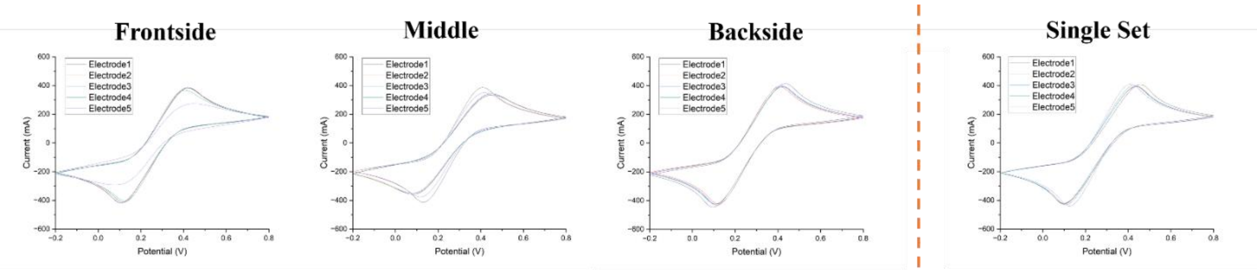


| (n=5) | |
|---|---|
| $i_{pa}/I_{pc}$ | 1.07 ± 0.013 |
| $\Delta E_p$ | 0.30 ± 0.020 |

| (n=5) | |
|---|---|
| $i_{pa}/I_{pc}$ | 1.07 ± 0.021 |
| $\Delta E_p$ | 0.32 ± 0.037 |

| (n=5) | |
|---|---|
| $i_{pa}/I_{pc}$ | 1.06 ± 0.002 |
| $\Delta E_p$ | 0.31 ± 0.017 |

| (n=5) | |
|---|---|
| $i_{pa}/I_{pc}$ | 1.05 ± 0.007 |
| $\Delta E_p$ | 0.32 ± 0.026 |

**Figure S2.** A) Position of precursor CCE arrays inside the pyrolysis oven. B) Electrochemical characterization of the CCE array chips pyrolyzed in different positions in the pyrolysis furnace: Comparison of three CCE chips pyrolyzed simultaneously with a single CCE array chip pyrolyzed alone. Cyclic voltammograms were generated in 10 mM hexacyanoferrate (II/III) at a scan rate of 100 $mV^{-s}$ with the potential range between -0.2 to 0.8V. The peak potential difference (ΔE) and the current peak height ratio between anodic ($i_{pa}$) and cathodic ($i_{pc}$) peaks are shown below each plot.

The resulting CVs and associated parameters (ΔE, $i_{pa}$, $i_{pc}$) show highly reproducible results verifying a highly reproducible fabrication process,

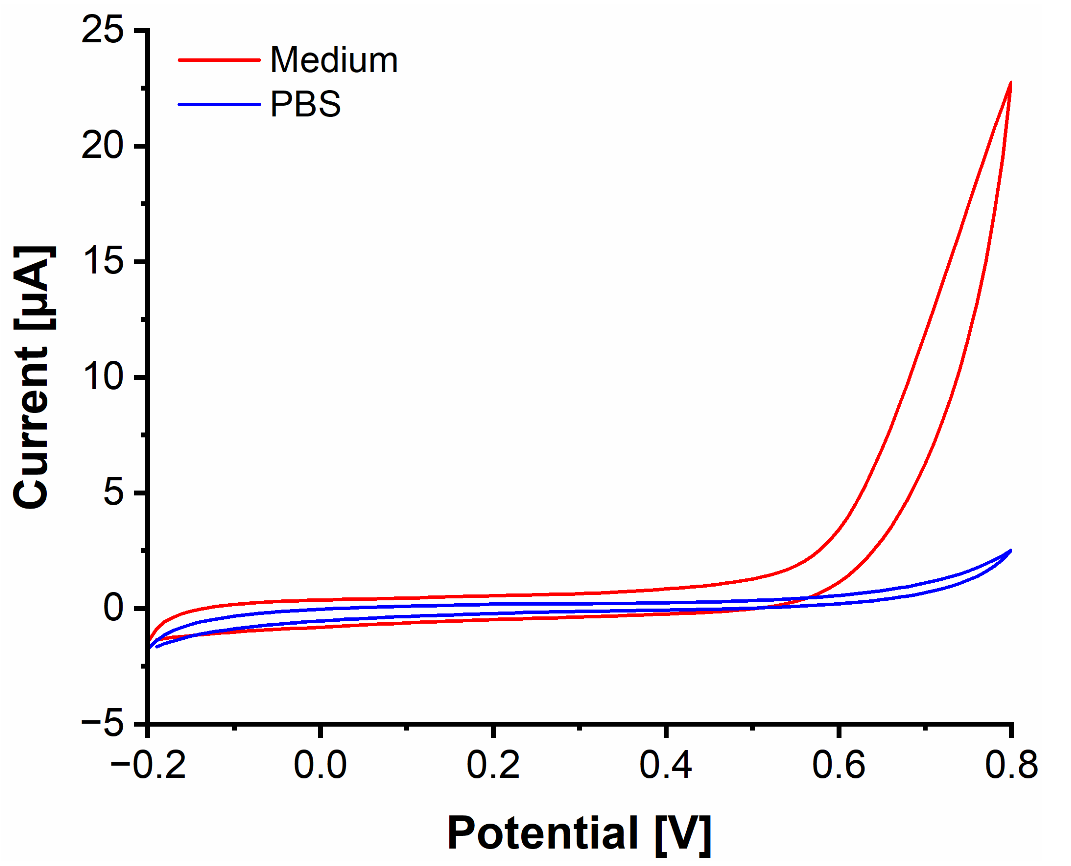


**Figure S3.** CVs generated in the neural basal media and PBS
